# An atomic interaction conserved for over 600 million years gates inhibitory neurotransmission

**DOI:** 10.64898/2026.05.22.727206

**Authors:** Cecilia M. Borghese, Netrang G. Desai, Pranavi Garlapati, Syedah K. Shah, Jason D. Galpin, Luthary Segura, Nandan Haloi, Harold H. Zakon, Rebecca J. Howard, Erik Lindahl, Christopher A. Ahern, Marcel P. Goldschen-Ohm

## Abstract

Pentameric ligand-gated ion channels (pLGICs) mediate fast inhibitory neurotransmission critical for neuronal network stability. A tyrosine residue in the M2-M3 linker of inhibitory pLGICs, conserved for over 600 million years, is positioned where it could hydrogen bond (H-bond) to the backbone of the neighboring Cys-loop. Given the pathogenic effects of variants of this tyrosine, we hypothesized that this H-bond stabilizes extracellular-to-transmembrane domain coupling essential for channel gating. To test this hypothesis, we used site-directed mutagenesis, noncanonical amino acid incorporation, and electrophysiological recordings in *Xenopus laevis* oocytes to disrupt this hydrogen bond in GABA_A_ and glycine receptors. Loss of this interaction via tyrosine substitutions or backbone amide modifications that ablate the acceptor or donor, respectively, markedly decrease agonist sensitivity and maximal channel activation, with effects localized to specific subunits. Molecular dynamics simulations support a role for this H-bond in channel gating. These findings reveal a critical atomic interaction underlying a shared mechanism for inhibitory receptor gating and provide a mechanistic explanation for disease-associated mutations linked to epilepsy, neurodevelopmental disability, and hyperekplexia.

## Introduction

Pentameric ligand-gated ion channels (pLGICs) are central to fast synaptic transmission and the dynamic regulation of neuronal excitability. pLGICs divide into excitatory, cation-selective receptors (e.g., nicotinic acetylcholine and 5-HT3) and inhibitory, anion-selective receptors (GABA_A_ and glycine) which counterbalance excitatory drive and stabilize network activity. GABA_A_ receptors (GABA_A_Rs) are typically heteromeric, commonly comprising combinations of α, β, and γ subunits, with subunit composition conferring distinct pharmacological and kinetic properties (Sallard et al., 2021). Glycine receptors (GlyRs) are also commonly heteromers comprised of four α subunits and one β subunit, with isoform-specific roles across developmental stages and circuits (Mizzi & Blundell, 2025). However, GlyRs can also form as α subunit homomers, a stoichiometry that has been widely studied in heterologous systems.

All pLGICs share a conserved modular organization: an extracellular domain (ECD) consisting largely of a β-sheet barrel connected to a four-helix transmembrane domain (TMD) bundle (M1–M4), with the pore formed primarily by the M2 helices from each subunit (**Fig. 1a**). Agonists (e.g., neurotransmitter) bind at orthosteric sites in the ECD at specific intersubunit interfaces distant from the main pore gate in the TMD. The transduction of the chemical energy from agonist-binding to gating of the ion pore involves several loops at the ECD–TMD interface including the β1-β2, β8-β9, and Cys loops, and the pre-M1 and M2-M3 linkers (**Fig. 1a**) (Michałowski et al., 2025).

**Figure 1.**
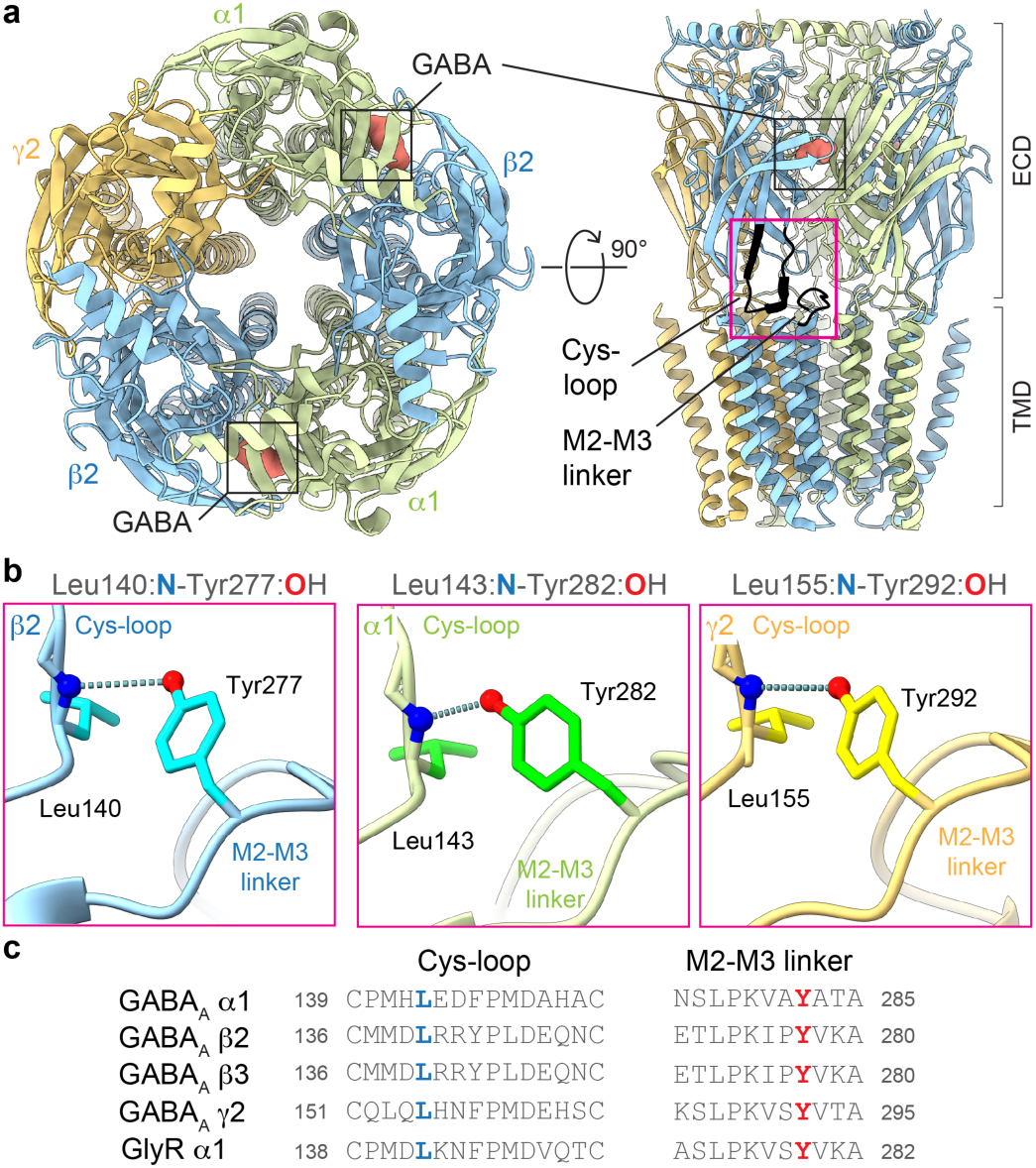
A predicted backbone-to-sidechain H-bond linking the Cys-loop to a conserved tyrosine in the M2-M3 linker. **a** Cryo-EM map (PDB 6X3Z) of human α1β2γ2 GABA_A_R with bound GABA (salmon) viewed from the side (left) and top (right) (ECD, extracellular domain; TMD, transmembrane domain). Cys-loop and M2-M3 linker colored black in one β2 subunit. **b** Predicted H-bonds β2(Leu140:N-Tyr277:OH) (left), α1(Leu143:N-Tyr282:OH) (center) and γ2(Leu155:N-Tyr292:OH) (right) shown as dashed lines. Structure visualizations with ChimeraX (Pettersen et al., 2004). **c** Alignment of the Cys-loop and M2-M3 linkers of subunits expressed in this study. Donor and acceptor residues involved in the H-bonds above are marked in bold.

Although decades of biophysical, structural, and electrophysiological work have delineated the overall architecture of pLGICs, precise molecular details of how ECD binding events are coupled to TMD gating remain incompletely understood. This gap is consequential as dysregulation of chloride conductance due to dysfunction of anionic-selective pLGICs is clinically implicated in neurodevelopmental and neuropsychiatric disorders including epilepsy, intellectual disability, Autism, schizophrenia, Alzheimer’s, and Fragile-X syndrome (Treviño et al., 2025), as well as movement disorders such as hyperekplexia (startle disease) (Mizzi & Blundell, 2025). These receptors are also major pharmacological targets for widely prescribed therapies to seizures, anxiety, insomnia, mood disorders, pain, muscle spasms, and alcohol withdrawal (e.g., benzodiazepines, Z-drugs, neurosteroids), as well as sedation (anesthetics) (Mizzi & Blundell, 2025; Treviño et al., 2025). Mechanistic insight into ECD–TMD coupling is therefore essential for interpreting pathogenic mutations and for rational drug design.

Here, we show that a specific hydrogen bond (H-bond) formed between the OH hydroxyl group of a highly conserved tyrosine sidechain in the M2-M3 linker and the main-chain amide nitrogen NH group of a leucine residue within the neighboring Cys-loop (**Fig. 1b,c, Supplementary Table 1**) is essential for efficient neurotransmitter-activation of both α1β2/3γ2 GABA_A_Rs and α1 GlyRs. Consistent with its functional importance, variants of the conserved tyrosine that disrupt this H-bond are associated with epilepsy and moderate to severe intellectual disability in β2/3 GABA_A_Rs subunits (Maillard et al., 2022; Mohammadi et al., 2024), or hyperekplexia in GlyRs (Shiang et al., 1995). We show that nearly identical functional deficits to channel activation are conferred by either 1) disruption of the acceptor OH group via substitution of the conserved tyrosine, or 2) disruption of the donor NH group by non-canonical α-hydroxy acid amide-to-ester substitution in the main-chain of the Cys-loop leucine. This strongly suggests that loss of this specific H-bond is the underlying mechanism for channel dysfunction in these pathogenic variants.

Cryo-EM structures of both heteromeric α1β2/3γ2 GABA_A_Rs (Kim et al., 2020; Sente et al., 2022) and homomeric α1 GlyRs (Yu et al., 2021) place the tyrosine OH group in proximity to the leucine amide nitrogen, consistent with an H-bond at this location in all subunits. However, we show that in heteromeric α1β2/3γ2 GABA_A_Rs this H-bond is only functionally relevant in β subunits and has no functional contribution in α1 or γ2 subunits. This aligns with both 1) the more severe effects of variants of the conserved tyrosine in β subunits and 2) our prior observations that perturbations in the M2-M3 linker of β2 subunits have differential effects from analogous perturbations in α1 or γ2 subunits (Borghese et al., 2025; Nors et al., 2024). Taken together, we hypothesize that β subunit gating-transduction loops have a unique role in channel function due to their location at the GABA-binding intersubunit interfaces.

Our findings delineate a specific, structurally-grounded interaction at the ECD–TMD interface that is necessary for robust inhibitory signaling in anionic-selective pLGICs. A phylogenetic analysis suggests this mechanism specifically evolved primarily in anion-selective and not cation-selective pLGICs of Bilateria and Cnidaria over 600 million years ago. This provides a mechanistic rationale for disease-causing mutations associated with epilepsy, neurodevelopmental disorders, and hyperekplexia and highlights subunit-dependent specialization relevant to therapeutic targeting.

## Results

### Disease-associated variants of a conserved tyrosine in the M2-M3 linker impair activation of a synaptic GABA_A_R

Previous studies have identified a highly conserved tyrosine residue in the M2-M3 linker of GABA_A_R β subunits as important for normal receptor function. For example, a loss-of-function (LOF) variant of this tyrosine, β2(Tyr277Cys), is associated with epilepsy and moderate to severe intellectual disability (reported as Tyr301Cys using HUGO amino acid numeration) (Mohammadi et al., 2024). Although the HUGO numeration, which starts numbering at the first methionine in the amino acid sequence, is more aligned with databases such as Uniprot (UniProt Consortium, 2025) and ClinVar (Landrum et al., 2018), it is common in the scientific literature to use a numeration that starts at the first residue in the mature protein following the signal peptide sequence as we do here.

We hypothesized that the LOF phenotype reflects impaired coupling between the M2-M3 linker and neighboring Cys-loop due to disruption of a backbone-to-sidechain H-bond, which in wild-type receptors is formed between the main-chain amide nitrogen of β2(Leu140) in the Cys-loop and the sidechain OH group of β2(Tyr277) in the M2-M3 linker (**Fig. 1b**). To test this hypothesis, we expressed α1β2γ2 GABA_A_Rs (or variants thereof) in *Xenopus laevis* oocytes and measured current responses to brief pulses of increasing concentrations of GABA (**Fig. 2a**). To estimate the maximal possible current in each cell, GABA concentration-response series were followed by co-application of saturating GABA plus 30 µM of the allosteric activator propofol (PPF) (**Fig. 2a**). Although it is likely that all channels are not open with exactly 100% probability during the peak response to co-application of GABA and PPF, normalizing currents to this maximal response nonetheless provides a useful estimate of the relative efficiency with which GABA alone can activate the channel (i.e., by comparison to maximal co-activation by GABA and PPF). Because PPF binds to the TMD and bypasses normal neurotransmitter ECD–TMD transduction (Kim et al., 2020), it is useful even for perturbations that disrupt this coupling. Finally, recordings were bookended by application of the pore blocker picrotoxin (PTX), which will block, and thus assay for, any spontaneous unliganded opening that may be conferred by mutations.

**Figure 2.**
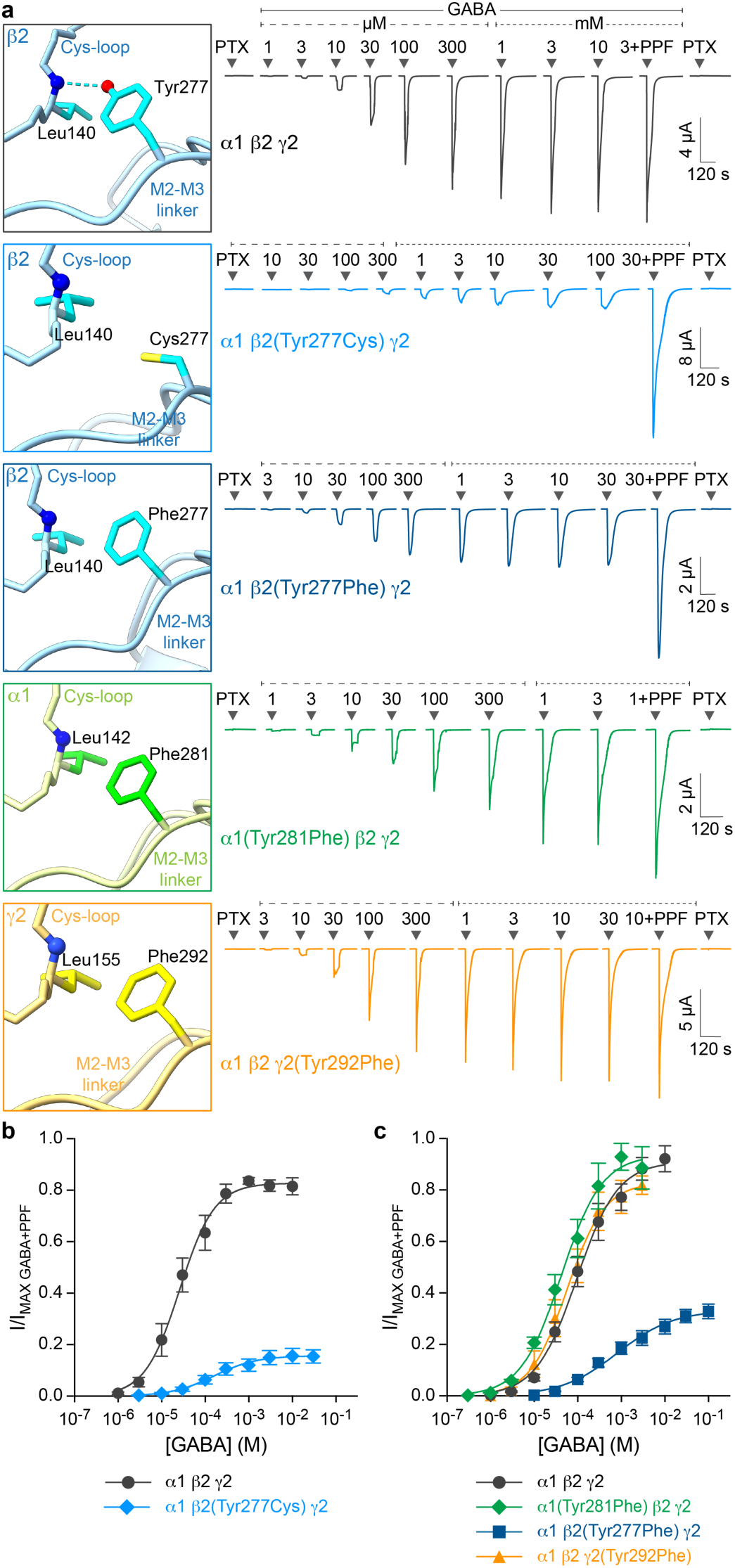
Substitutions of a conserved tyrosine in the M2-M3 linker impair channel gating specifically in β subunits. **a** AlphaFold structures (left) and representative current traces (right) for wild-type and M2-M3 tyrosine substitutions in GABA_A_R subunits. Arrows above traces indicate approximate onset of pulses of either 1 mM picrotoxin (PTX), increasing concentrations of GABA, or maximal GABA + 30 µM propofol (PPF). **b**,**c** Concentration-response relations for GABA-elicited currents, normalized to maximal GABA + PPF responses. Data are mean ± SEM across oocytes. Curves are fits of the Hill equation (Eq. 1) to the means. See Supplementary Table 3 for summary statistics and fit parameters. Number of oocytes (n) are: α1β2γ2: n = 5 or 6*; α1β2(Tyr277Cys)γ2: n = 7; α1β2γ2: n = 6; α1(Tyr281Phe)β2γ2: n = 5; α1β2(Tyr277Phe)γ2: n = 10; α1β2γ2(Tyr292Phe): n = 5. Note: The cysteine substitution was studied in human receptors, whereas phenylalanine substitutions were studied in rat receptors, which are for our purposes are essentially identical. One subtle difference is that the amino acid numeration differs by one between rat and human α1 subunits (e.g., compare with human numeration in Fig. 1). *Numbers for both human and rat wild-type receptors for appropriate comparison with mutants.

We show that α1β2(Tyr277Cys)γ2 receptors are not only less sensitive to GABA (i.e., have right-shifted GABA concentration-response curves), but also are maximally opened by GABA with much lower efficiency than wild-type receptors (**Fig. 2a,b**). Thus, GABA acts as a weak partial agonist at α1β2(Tyr277Cys)γ2 receptors, where saturating GABA can only elicit ∼20% of the maximal possible channel current (i.e., as measured by co-applying GABA and PPF) as opposed to the ∼80% activation observed for wild-type receptors. These results are consistent with previous observations for the LOF phenotype in α1β2(Tyr277Cys)γ2 receptors (Mohammadi et al., 2024). As expected for a LOF mutation, no PTX-sensitive unliganded opening was observed.

Another variant of this conserved tyrosine, β2(Tyr277Phe) is linked to intellectual disability and epileptic encephalopathy (**Supplementary Table 2**). This tyrosine to phenylalanine substitution specifically removes the H-bond acceptor OH group with little other changes to the sidechain (**Fig. 2a**). If loss of the H-bond is the underlying mechanism for dysfunction of both β2(Tyr277Cys) and β2(Tyr277Phe) variants, we expect they should confer similar functional deficits. Consistent with this idea, we observe a similar loss of both GABA-sensitivity and maximal efficiency of channel opening for α1β2(Tyr277Phe)γ2 receptors (**Fig. 2a,c**).

### A specific H-bond linking neighboring gating-transduction loops

To further substantiate that the observed dysfunction conferred by substitutions of β2(Tyr277) are due to disruption of the H-bond β2(Leu140:N-Tyr277:OH) (**Fig. 3a,c**) and not simply differences in sidechains, we specifically removed the donor amide nitrogen using an amide-to-ester substitution in the main-chain of β2(Leu140). Because conventional amino acid substitutions do not alter the composition of the main-chain, we used a noncanonical amino acid (ncAA). Specifically, we substituted β2(Leu140) with the cognate α-hydroxy acid for leucine (Lah) to swap the amide peptide bond for an ester bond without altering the sidechain (**Fig. 3a**). This specifically replaces the backbone amide nitrogen with an oxygen which disrupts the H-bond donor in an analogous fashion to disruption of the H-bond acceptor by substitutions of the β2(Tyr277) sidechain that ablate the OH group (**Fig. 2a**).

**Figure 3.**
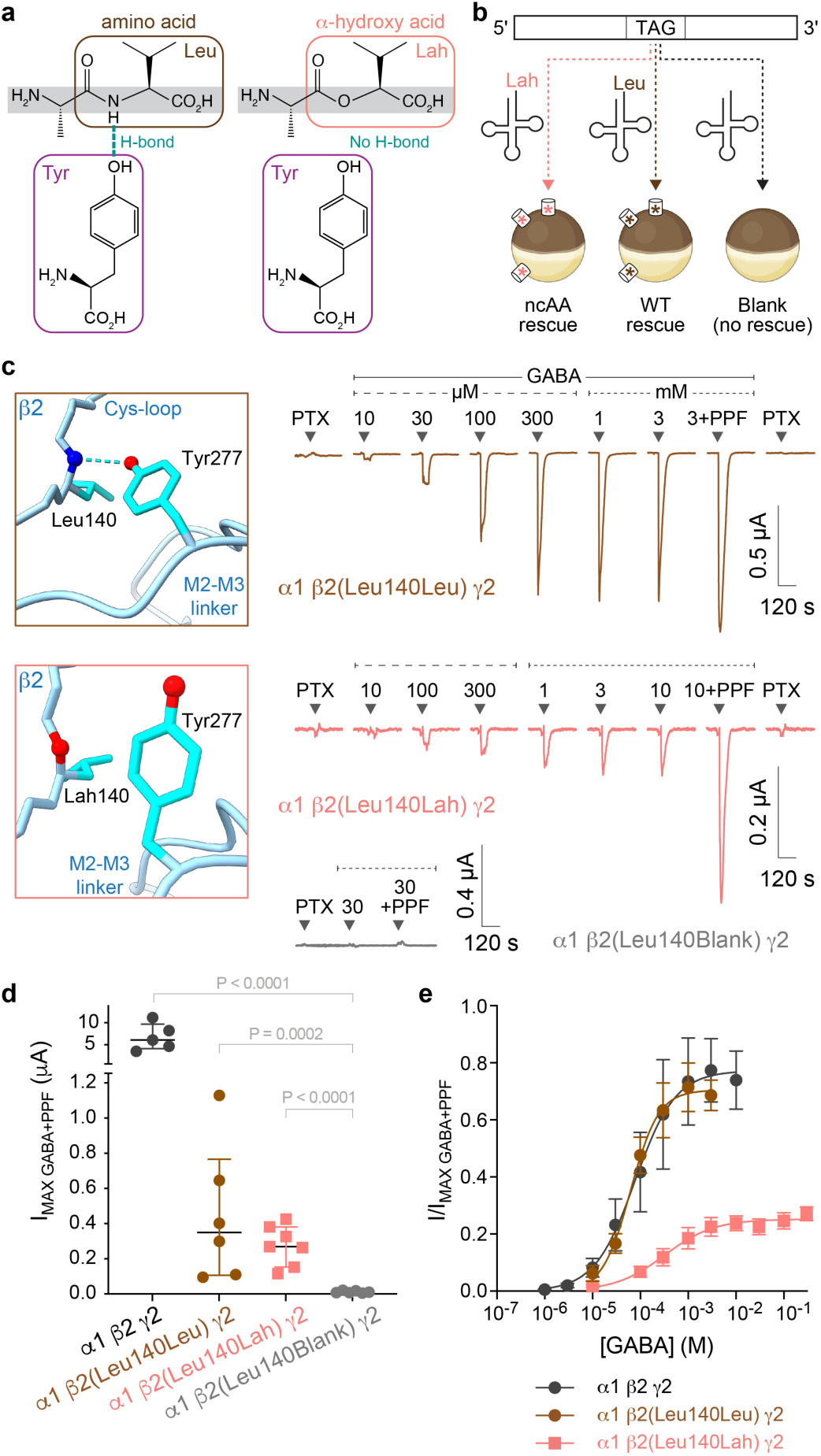
A backbone-to-sidechain H-bond linking the Cys-loop and M2-M3 linker is crucial for robust channel activation. **a** Chemical structures showing *left*) an H-bond between a backbone amide nitrogen of a leucine in the Cys-loop and the OH group of a tyrosine in the M2-M3 linker, and *right*) loss of this H-bond capacity upon substitution of the leucine with its cognate α-hydroxy acid (Lah) which confers an amide-to-ester swap in the Cys-loop main-chain without altering the sidechain. **b** Schematic of the in vivo nonsense suppression incorporation of α-hydroxy acids and controls (see main text). **c** AlphaFold structures (left) and representative current traces (right) for GABA_A_R wild-type rescue α1β2(Leu140Leu)γ2, amide-to-ester swap α1β2(Leu140Lah)γ2, and read-through control α1β2(Leu140Blank)γ2. Arrows above traces indicate approximate onset of pulses of either 1 mM picrotoxin (PTX), increasing concentrations of GABA, or maximal GABA + 30 µM propofol (PPF). **d** Total current per oocyte obtained in the presence of maximal GABA + 30 µM propofol suggests reliable nonsense suppression incorporation of Leu and Lah with little to no read-through (i.e., relative lack of current for Blank). Number of oocytes for each condition (left to right) are n = 5, 6, 7, 6. Plots show median and interquartile intervals. P-values from Kruskal-Wallis ANOVA with posthoc Dunn’s multiple comparisons test are indicated. **e** Concentration-response relations for GABA-elicited currents, normalized to the response to maximal GABA + PPF. Data are mean ± SEM across oocytes. Curves are fits of the Hill equation (Eq. 1) to the means. See Supplementary Table 3 for summary statistics and fit parameters. Number of oocytes (n) are: α1β2γ2: n = 5; α1β2(Leu140Leu)γ2: n = 6; α1β2(Leu140Lah)γ2: n = 7.

We used in vivo nonsense suppression to introduce the α-hydroxy acid Lah (**Fig. 3a**) as previously described (Borghese et al., 2025; Leisle et al., 2022). Briefly, the codon encoding the residue of interest is substituted with a TAG stop codon in the cDNA. The cRNAs encoding the subunits are injected along with an orthogonal tRNA directed to the UAG stop codon and ligated to either the wild-type amino acid (e.g., Leu), the α-hydroxy acid (e.g., Lah), or nothing (negative control, Blank) (**Fig. 3b**). The first condition is a control for nonsense suppression insertion at the site of interest, which if successful will result in wild-type protein. The second condition is the ncAA substitution. The last condition is a control to check for read-through which should result in no expression if translation stops at the UAG codon as expected. The absence of clear GABA+PPF-elicited currents when oocytes were co-injected with tRNA-Blank indicates there was little to no read-through, whereas nonsense suppression incorporation of Leu or Lah resulted in obvious GABA-elicited currents (**Fig. 3c,d**). Nonsense suppression incorporation of the wild-type amino acid Leu largely recapitulated wild-type function, indicating successful site-specific incorporation of synthesized amino acids (**Fig. 3c,e**). In contrast, incorporation of Lah resulted in a marked decrease in both GABA-sensitivity and the efficiency of channel opening by GABA (**Fig. 3c,e**). The analogous outcomes of disrupting the H-bond donor (amide-to-ester substitution in the Cys-loop main-chain) or acceptor (sidechain substitutions of the conserved M2-M3 tyrosine) strongly suggest that the H-bond β2(Leu140:N-Tyr277:OH) is essential for normal channel activation by GABA. And furthermore, that functional deficits conferred by variants of the conserved tyrosine β2(Tyr277) are due to disruption of this H-bond.

### Despite structural homology in all subunits, this specific loop-linkage is only functionally relevant in β subunits

The M2-M3 linker tyrosine discussed above is conserved in nearly all GABA_A_R subunits, except for δ, π and θ (**Supplementary Table 1**). Cryo-EM structural models further suggest that a homologous H-bond linking this conserved tyrosine to the backbone of the neighboring Cys-loop exists in α1 and γ2 subunits. To test whether channel gating depends on this linkage in other subunits, we explored tyrosine-to-phenylalanine substitutions that disrupt the H-bond acceptor in α1 or γ2 subunits (**Fig. 2a**). In contrast to the LOF effects observed for α1β2(Tyr277Phe)γ2 receptors, neither α1(Tyr281Phe)β2γ2 nor α1β2γ2(Tyr292Phe) had any effect on channel function (**Fig. 2c**). Thus, regardless of whether a homologous H-bond occurs in α1 or γ2 subunits, it is irrelevant to channel function.

A variant of the homologous conserved tyrosine in the β3 subunit, β3(Tyr277Cys) (reported as Tyr302Cys using HUGO amino acid numeration), has been characterized as LOF and is associated with epilepsy and intellectual disability. This variant confers a reduction in GABA efficacy (Absalom et al., 2019, 2022; Møller et al., 2017) and altered kinetics along with reduced GABA-induced current density, but normal expression levels in the plasma membrane (Janve et al., 2016). Given that synaptic GABA_A_Rs are often formed as heteropentamers comprised of α1, β1-3, and γ2 subunits (Sallard et al., 2021), we hypothesize that this H-bond has a similar functional role in all β subunits. To test this idea, we observed the functional effects of disrupting either the donor or acceptor in the β3 subunit in an analogous fashion to our approach in the β2 subunit. We show that either ablation of the acceptor OH group in α1β3(Tyr277Cys)γ2 or α1β3(Tyr277Phe)γ2 receptors, or ablation of the donor amide nitrogen in α1β3(Leu277Lah)γ2 receptors, all confer similar LOF effects analogous to those observed upon disruption of the homologous H-bond in the β2 subunit (**Supplementary Fig. 1**). This strongly suggests that normal channel activation by GABA requires this H-bond in whichever β subunit is present in the channel.

### H-bond dynamics for state-dependent coupling between gating-loops

To investigate the local structural ramifications of this loop linkage, we carried out all-atom molecular dynamics (MD) simulations starting from either closed (antagonist-bound) or desensitized (GABA-bound) cryo-EM structures of α1β2γ2 GABA_A_ receptors, or a simulated open-state structure (Haloi et al., 2025). To explore the local effects of ablating the H-bond discussed above, we additionally carried out MD simulations with the ncAA substitution β2(Leu140Lah). Given the importance of the H-bond linking the Cys-loop and M2-M3 linker in channel activation, we reasoned that any structural effects upon its ablation may be most prominent in an activated conformation. Thus, the ncAA substitution was introduced into the desensitized structure.

Donor-acceptor distance distributions for the Leu:N-Tyr:OH H-bond linking the Cys-loop and M2-M3 linker in each subunit suggest that this H-bond is highly stable in α1 subunits independent of conformation (**Fig. 4, Supplementary Fig. 2**). In contrast, the homologous H-bond in β2 and γ2 subunits are more dynamic, fluctuating between proximal (e.g., bonded) and distant (e.g., non-bonded) orientations. This fluctuation is reduced in the open conformation where the H-bond is primarily intact in β2 subunits and absent in the γ2 subunit, suggesting some degree of state-dependence to this bond in non-α subunits, and consistent with its importance in β2 subunits for channel opening. Ablation of the H-bond donor with the β2(Leu140Lah) amide-to-ester ncAA substitution conferred a much higher probability of the Tyr:OH to adopt an orientation distant from the Cys-loop, consistent with the idea that this substitution disrupts this particular linkage between the Cys-loop and M2-M3 linker. Although β2(Leu140Lah) had no effect on the homologous H-bond in α1 subunits, it is correlated with a stabilized H-bond in the γ2 subunit, the overall effects of which are consistent with changes in this loop linkage associated with a non-open conformation. Interestingly, differences in H-bond stability between conducting and non-conducting conformations were similar for both closed and desensitized states. Thus, it is possible that the identified H-bond is specifically stabilized in an asymmetric open conformation but becomes more dynamic in relatively symmetric non-conducting conformations (Mihaylov et al., 2025).

**Figure 4.**
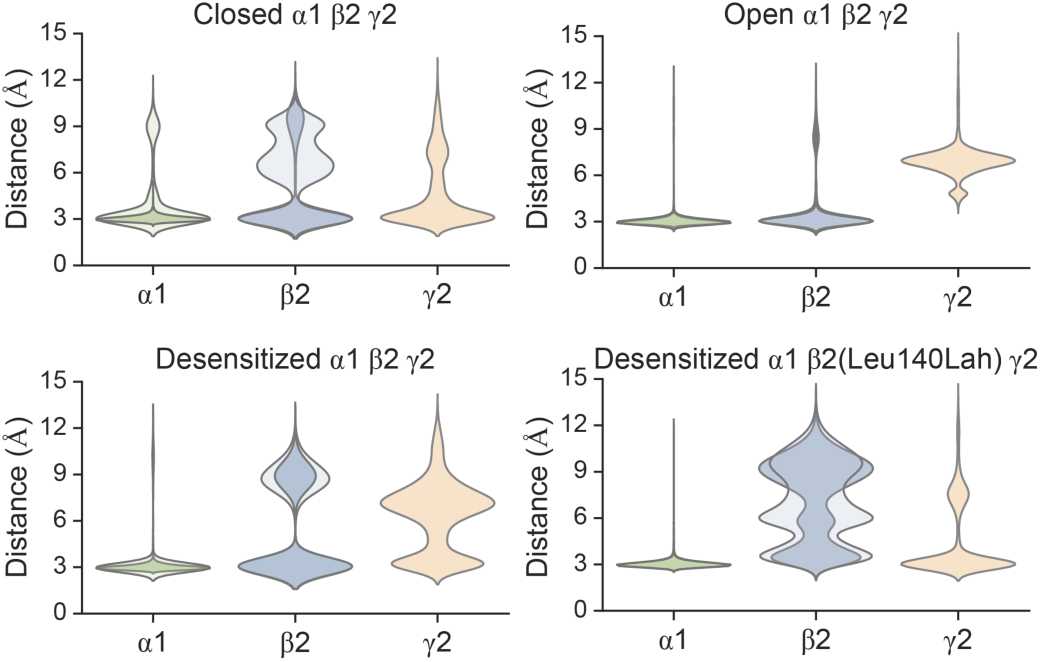
Dynamics of an H-bond linking neighboring gating loops. Donor-acceptor distance distributions from MD simulations of α1β2γ2 GABA_A_Rs in closed (antagonist-bound, PDB 6X3S), open (simulated structure (Haloi et al., 2025)), and desensitized (GABA-bound, PDB 6X3Z) states with and without the β2(Leu140Lah) amide-to-ester ncAA substitution. Distances are between a leucine backbone amide nitrogen in the Cys-loop (or the ester oxygen for the ncAA substitution) and a tyrosine OH group in the M2-M3 linker of the same subunit. Residue pairs are α1(Leu143:N-Tyr282:OH), β2(Leu140:N-Tyr277:OH), γ2(Leu155:N-Tyr292:OH), or β2(Lah140:O-Tyr277:OH). Distributions are aggregates over all four replicates, and distinct chains of the same subunit are overlaid (darker: chains A-B, lighter: chains C-D).

The backbone geometry of the ester peptide bond for the β2(Leu140Lah) substitution was similar to that of the wild-type amide peptide bond in the MD simulations of the desensitized state (**Supplementary Figs. 3-4**). However, the angular fluctuations were larger for the ester bond, indicating a more flexible backbone primarily localized to the substitution site, but with some additional flexibility in the immediately adjacent amide bonds as well. Although the ester peptide bond may confer increased backbone flexibility, the similar functional effects of the amide-to-ester swap in the Cys-loop and tyrosine substitutions in the M2-M3 linker suggest that loss of the H-bond linking these gating loops is the primary dysfunction. Furthermore, loss of this H-bond stabilization could also explain the localized increase in backbone flexibility of the Cys-loop.

Overall, these simulations suggest some state-dependence to the dynamics of this H-bond linkage between the Cys-loop and M2-M3 linker within β2 and γ2, but not α1 subunits. The open-state stabilization of the H-bond primarily in β2 is largely consistent with our observations for its functional importance only in these subunits. Nonetheless, the overall variation in these simulations suggests they be interpreted as a cautious suggestion rather than strong evidence.

### A conserved mechanism in anionic-selective GABA_A_ and glycine receptors

Anionic chloride-selective pLGICs include both GABA_A_ and glycine receptors. A homologous M2-M3 linker tyrosine is also conserved in glycine receptor subunits, and variants of this tyrosine are linked to hyperekplexia (startle disease) (Shiang et al., 1995) (**Supplementary Tables 1-2**). Cryo-EM structural models of GlyRs place the tyrosine’s OH group in proximity to a backbone amide nitrogen in the neighboring Cys-loop analogous to that observed in GABA_A_Rs (**Fig. 5a**). Thus, we hypothesized that both anionic receptors may share a similar mechanism in coupling neurotransmitter binding to channel gating involving a homologous H-bond linking neighboring transduction loops. To test this idea, we expressed GlyR α1 homomers in *Xenopus laevis* oocytes and measured functional activation by glycine. Similar to GABA_A_Rs, we assessed maximal channel activation using co-application of glycine and the allosteric activator PPF and assayed for unliganded channel opening by applying the pore-blocker PTX (**Fig. 5a**).

**Figure 5.**
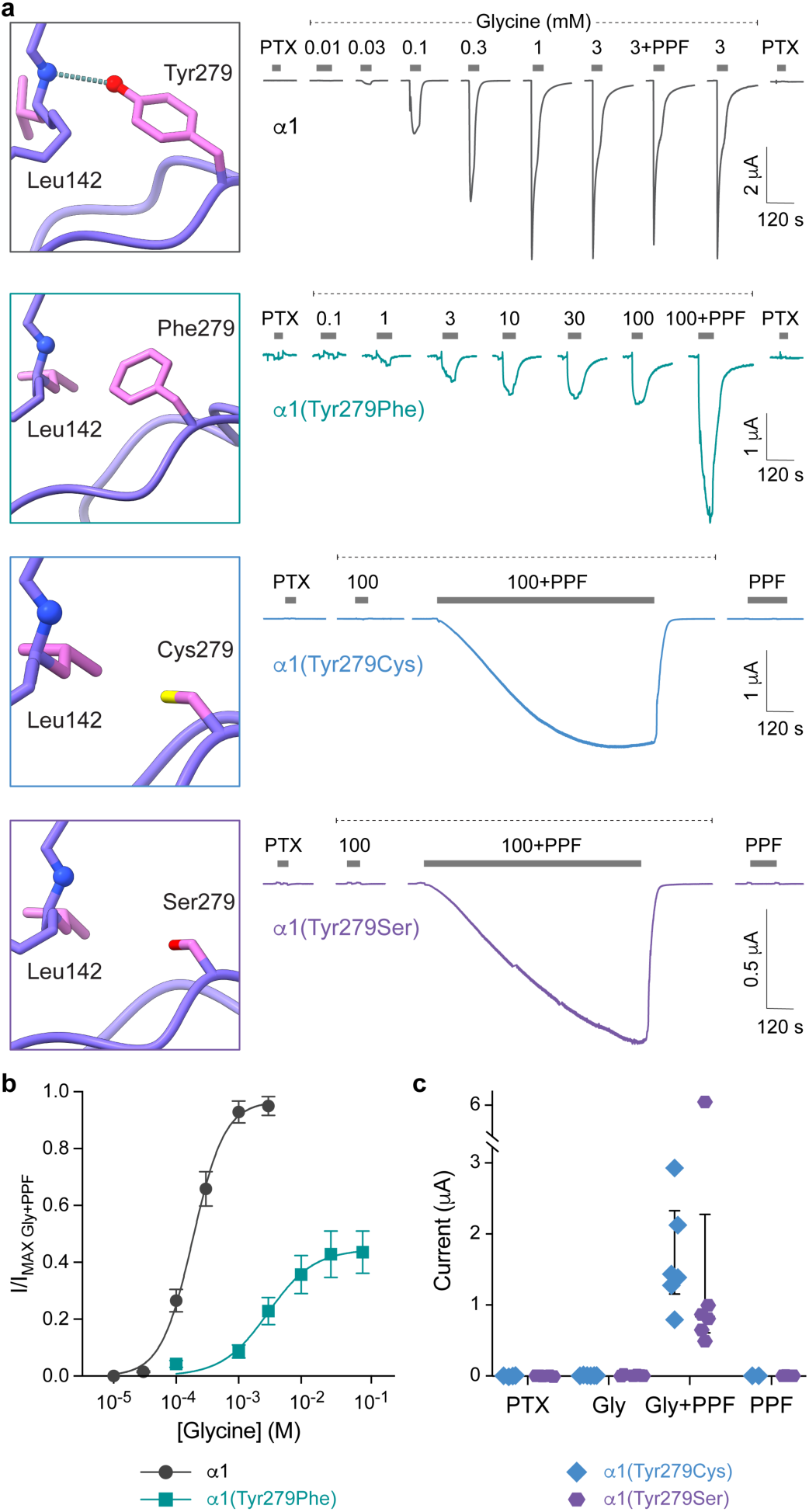
A conserved mechanism in glycine receptors. **a** Cryo-EM map (PDB 6PM3) or AlphaFold structures (left) and representative current traces (right) for wild-type and M2-M3 tyrosine substitutions in α1 GlyRs. Bars above traces indicate approximate application of pulses of either 1 mM picrotoxin (PTX), glycine, or maximal glycine + 100 µM propofol (PPF). **b** Concentration-response relations for glycine-elicited currents, normalized to the response to either maximal glycine or maximal glycine + PPF, whichever is greater. Data are mean ± SEM across oocytes. Curves are fits of the Hill equation (Eq. 1) to the means. See Supplementary Table 3 for summary statistics and fit parameters. Number of oocytes (n) are: α1: n = 6; α1(Tyr279Phe): n = 6. **c** Current amplitudes from α1(Tyr279Cys) and α1(Tyr277Ser) GlyRs evoked by different ligands. Number of oocytes (n) per condition (from left to right) are n = 4, 6; 6, 2; 5, 6; 6, 4.

Ablating the H-bond acceptor OH group via substitution of the conserved tyrosine with phenylalanine resulted in a similar LOF phenotype to that observed upon H-bond disruption in GABA_A_Rs. Namely, GlyR α1(Tyr279Phe) receptors have both a reduced glycine sensitivity and much lower maximal efficiency of channel opening as compared to wild-type GlyR α1 receptors (**Fig. 5a,b**). This mutant also exhibits visually slower activation kinetics, although we did not attempt to quantify this as precise kinetic measurements are challenged in oocytes due to slow solution exchange over the large cell surface and highly invaginated membrane. As expected for LOF, no obvious PTX-sensitive current indicative of unliganded opening was observed.

Known pathogenic variants of this conserved tyrosine include substitutions with either cysteine or serine, both of which are linked to hyperekplexia (Shiang et al., 1995) (**Supplementary Table 2**). We were unable to elicit current responses to glycine in either α1(Tyr279Cys) or α1(Tyr279Ser) GlyRs, nor did these channels respond to either PTX or PPF alone (**Fig. 5a,c**). However, current responses from both α1(Tyr279Cys) and α1(Tyr279Ser) GlyRs were evoked upon co-application of glycine and PPF, indicating that the channels did express and were functional. These currents activated with much slower kinetics as compared to glycine-evoked responses in wild-type receptors, consistent with highly impaired gating in these mutants. The more severe LOF effects for cysteine or serine substitutions of the conserved tyrosine in GlyRs as compared to cysteine substitution in GABA_A_Rs could reflect a more pronounced disabling effect when these substitutions are present in homomeric receptors as opposed to only β subunits within heteromeric GABA_A_Rs. Our results are largely consistent with prior observations for LOF effects in α1(Tyr279Cys) GlyRs, including decreased sensitivity to glycine and reduced partial agonists efficacies when expressed in HEK293 cells (Lynch et al., 1997). Why the LOF is more severe in oocytes (e.g., no activation by glycine alone) is unclear.

We also attempted to disrupt the H-bond donor using an α-hydroxy acid amide-to-ester substitution at α1(Leu142) in the Cys-loop main-chain. However, whereas nonsense suppression rescue of wild-type GlyRs resulted in normal glycine-evoked currents, no currents could be evoked with glycine and/or PPF upon incorporation of the cognate α-hydroxy acid Lah. It is unclear whether α1(Leu142Lah) GlyRs are non-functional or do not express. Either way, our observations strongly support an essential role for this H-bond in glycine receptor activation, similar to that for GABA_A_Rs.

### Evolutionarily conserved for over 600 million years

We collected a dataset consisting of protein sequences for pLGICs from representative organisms of diverse eukaryotes spanning unicellular and multicellular organisms from algae to humans (**Supplementary Table 4**). We limited our investigation to eukaryotes as a more inclusive study that also included prokaryotes and other groups showed that eukaryotic pLGICs are monophyletic (Jaiteh et al., 2016). The sequences were aligned and identified as either anion- or cation-selective channels based on widely accepted permeability motifs in the M2 α-helix (Nemecz et al., 2016) (see methods). Sequences with ambiguous ion-selectivity were not further considered. Those sequences with non-ambiguous ion-selectivity were classified based on whether they contained a tyrosine at the homologous position with the conserved tyrosine in the M2-M3 linker discussed above. Our time tree (Kumar et al., 2022) of eukaryotic organisms reveals that tyrosine at this position in the M2-M3 linker is phylogenetically ancient in most metazoan (animal) anion-selective pLGICs but is nearly absent in cation-selective channels (**Fig. 6**). The full phylogenetic tree is available in **Supplementary File 1** and at https://itol.embl.de/shared/1ieszf2NJt15U. Cnidaria (*Nematostella vectensis* and *Hydra vulgaris*) and Bilateria (*Homo sapiens*, *Drosophila melanogaster* and *Caenorhabditis elegans*) subunits can be largely identified as forming anion- or cation-selective channels based solely on whether a tyrosine exists in this position, suggesting a very strong selection pressure. Of the human sequences, the few exceptions are three subunits that do not supply the principal component to the orthosteric binding site (δ, π and θ), as do GABA_A_R β and GlyR α subunits, where we identified a functional importance of this residue.

**Figure 6.**
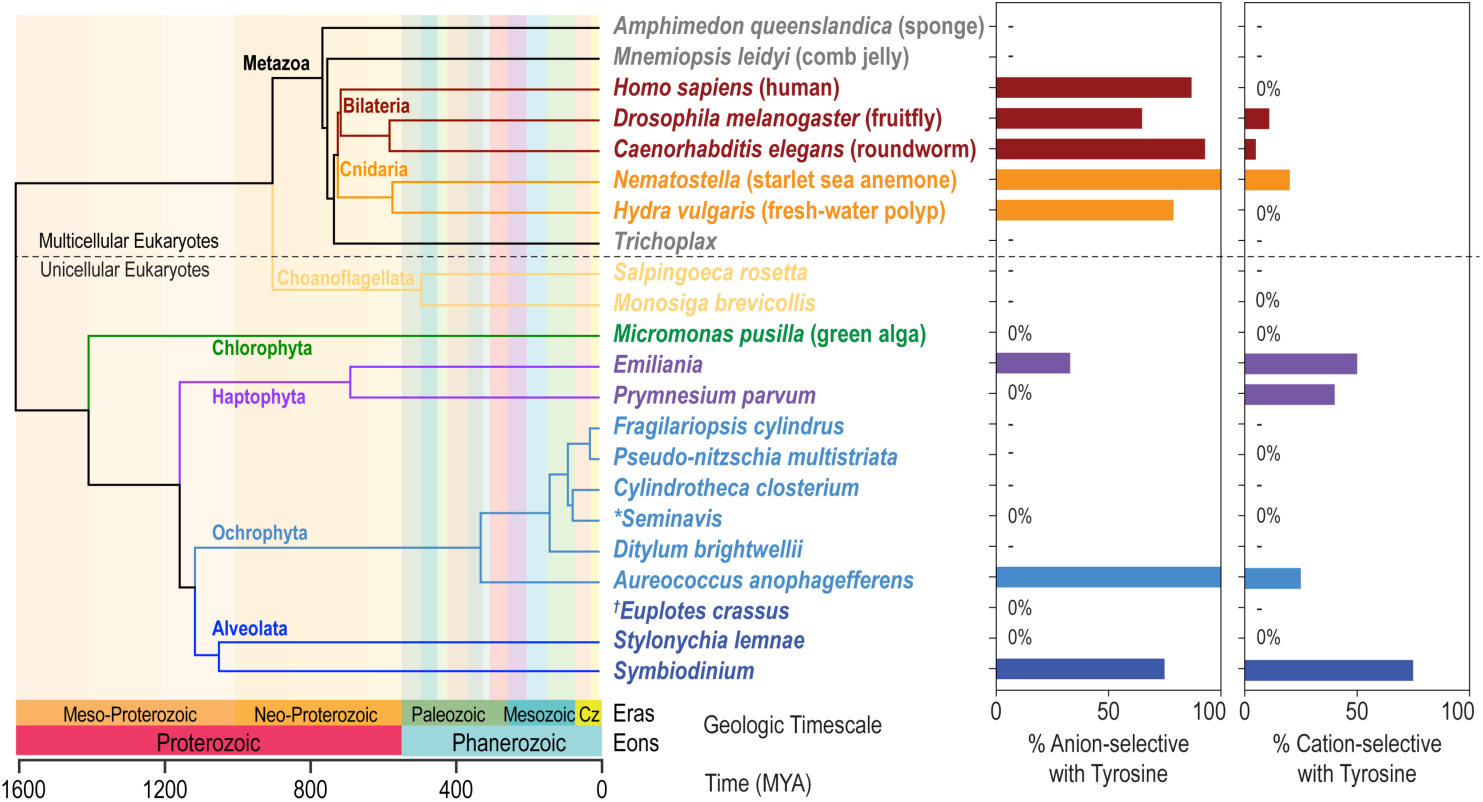
Evolutionary perspective for a tyrosine in the M2-M3 linker of anionic and cationic pLGICs. Time tree of representative eukaryotic organisms included in the phylogenetic analysis (**Supplementary Table 4**; **Supplementary File 1**). For each organism the percentage of anion-and cation-selective pLGIC sequences that contain a tyrosine in the M2-M3 linker at the homologous position with GABA_A_ β2(Tyr277) are shown to the right. Dashes indicate that there are no identifiable anionic or cationic pLGIC for that organism. \**Seminavis robusta* replaced with *Navicula veneta*. ^†^*Euplotes crassus* was not available in the database but is closely related to *Stylonychia lemnae*.

In other metazoans (Placozoa, Porifera and Ctenophora), with contentious phylogenetic relationships to Cnidaria and Bilateria, we did not detect any pLGICs. A more exhaustive search of the genomes of several Porifera and Ctenophora species also failed to find any pLGICs in a recent study (Ojha et al., 2026). Interestingly, Porifera and Placozoa do not have a nervous system, and Ctenophora lacks a central nervous system (Burkhardt et al., 2023), which may explain the absence of clear pLGIC homologs in these organisms.

The correlation between the presence of this tyrosine and the channel selectivity is not present in protist (unicellular eukaryotes) (**Fig. 6**). Although some protists have a tyrosine in this position, acquired independently, via lateral gene transfer, or from an ancestral receptor, many others do not irrespective of the inferred channel selectivity. We hypothesize that there are very different selection pressures on these channels in multicellular organisms with nervous systems versus unicellular organisms, which has contributed to the segregation of this specific loop-linkage mechanism for anion-selective pLGICs in many metazoans. In contrast, these same multicellular organisms appear to have evolved a distinctly different mechanism for cation-selective channels for which there is a variety of non-tyrosine residues at this position. An exception is a more ancient outgroup subfamily of cationic channels from primarily unicellular organisms that mostly have a tyrosine here which likely evolved independently (**Supplementary File 1**).

We conclude that in more complex animals that possess a nervous system (Cnidaria and Bilateria), most anion-selective pLGICs evolved a distinct H-bond interaction essential to robust neurotransmitter-activation over 600 million years ago, a distinct mechanism that is largely absent in cation-selective channels.

## Discussion

Here we show that a main-chain to sidechain H-bond linking the Cys-loop and neighboring M2-M3 linker is crucial for normal neurotransmitter binding-to-gating transduction in both GABA_A_ and glycine receptors. Disruption of this H-bond results in both loss of neurotransmitter-sensitivity and a drastic reduction in maximal neurotransmitter-evoked currents which we interpret as a reduction in the efficiency with which neurotransmitter can open the channel. Although this H-bond appears to be structurally conserved in all subunits of heteromeric α1β2-3γ2 GABA_A_Rs, it is only functionally relevant in β subunits. We hypothesize that this H-bond only contributes to gating within subunits that provide the principal face of the orthosteric neurotransmitter binding site and thus place their Cys-loop and M2-M3 linker directly below the neurotransmitter binding site. For example, GABA binds at β-α inter-subunit interfaces, with β providing the principal face of the binding site which is located directly above the β subunit Cys-loop and M2-M3 linker. This hypothesis is consistent with our recent findings of differential effects for M2-M3 linker perturbations in β vs non-β subunits (Borghese et al., 2025; Nors et al., 2021, 2024). Furthermore, LOF due to disruption of this H-bond accounts for a mechanism by which pathogenic variants of the conserved M2-M3 linker tyrosine cause epilepsy, neurodevelopmental disability, or hyperekplexia (startle disease). This knowledge enables future development of targeted therapies aimed at either re-linking the lost connection between transduction loops or modulating this interaction in wild-type channels to regulate activity.

The high conservation of the M2-M3 tyrosine in anion-selective inhibitory pLGICs underscores its importance (**Fig. 6**, **Supplementary Tables 1, 4**). Replacing this tyrosine in GABA_A_R β2/3 or GlyR α1 subunits with substitutions linked to epilepsy and neurodevelopmental disability or hyperekplexia results in highly similar functional deficits. These observations strongly suggest that this tyrosine is involved in an essential H-bond that is part of a shared gating mechanism amongst inhibitory pLGICs. Interestingly, cationic-selective pLGICs in human and other chordate organisms lack a tyrosine in the homologous position of their M2-M3 linkers (**Supplementary Table 1**), and analysis of the corresponding structures does not reveal an analogous H-bond. Thus, despite overall homology suggesting that the gross gating motions are similar in all pLGICs (Cymes & Grosman, 2021), this specific mechanism has evolved primarily in anion- and not cation-selective pLGICs.

In summary, we demonstrate using traditional and non-canonical residue substitutions that a hydrogen bond between the M2-M3 linker and the Cys-loop is a pivotal determinant of ECD–TMD coupling in inhibitory pLGICs. We further connect clinically relevant mutations to a specific mechanistic breakdown, showing how the loss of a single atomic interaction can reduce both agonist sensitivity and gating efficiency. In addition, we uncover another example of subunit asymmetry in heteromeric GABA_A_Rs, with implications for designing subunit-selective modulators and for interpreting genetic variation across patient populations. By situating this precise molecular contact within the broader structural and physiological framework of inhibitory receptors, our study offers both mechanistic clarity and translational relevance.

## Methods

All procedures are essentially the same as in Borghese et al., 2025 (Borghese et al., 2025).

### Mutagenesis and in vitro transcription

DNA for wild-type and mutant GABA_A_R rat and human α1, β2, β3, and γ2 subunits was subcloned in the pUNIV vector (Venkatachalan et al., 2007). The mature protein numeration for β2 and γ2 subunits is the same for rat and human, but the numeration for the rat α1 subunit is (human numeration -1) for most of the subunit. DNA for wild-type and mutant GlyR human α1 subunit was subcloned in the pBK-CMV vector. For ncAA incorporation, the codon of the residue of interest was replaced by the TAG stop codon (in contrast to TGA stop codons for each subunit). Mutations were introduced by GenScript and confirmed by sequencing of the entire subunit. Complementary RNA (cRNA) for each construct was generated (mMessage mMachine T7, Ambion) and quantified (Qubit, ThermoFisher Scientific) prior to injection in *Xenopus laevis* oocytes.

### ncAA synthesis

For nonsense suppression in GABA_A_ subunits, we used TAG mutants of the β2 and β3 GABA_A_ subunits and PylT tRNAs in *Xenopus laevis* oocytes. PylT lacking the two terminal CA nucleotides was synthetized by Integrated DNA Technologies, Inc., folded and misacylated as previously described (Infield et al., 2018). Leu- and α-hydroxy Leu- (Lah) pdCpA-substrates were synthesized according to published procedures (Infield et al., 2018).

### TEVC recording in oocytes

Defolliculated *Xenopus laevis* oocytes were obtained from Ecocyte. Oocytes were injected with 12 ng of total cRNA for GABA_A_ α1, β2, and γ2 subunits (wild-type or mutants) in a 1:1:10 ratio (Boileau et al., 2002), or with 5-24 ng of cRNA for GlyR α1 (Nanoject, Drummond Scientific). When injecting the TAG mutant of a subunit, 125 ng of tRNA was mixed with the cRNA encoding the GABA_A_ subunits or GlyR α1. Oocytes were incubated in a sterile incubation solution (88 mM NaCl, 1mM KCl, 2.4 mM NaHCO_3_, 19 mM HEPES, 0.82 mM MgSO_4_, 0.33 mM Ca(NO_3_)_2_, 0.91 mM CaCl_2_, 10,000 units/L penicillin, 50 mg/L gentamicin, 90 mg/L theophylline, and 220 mg/L sodium pyruvate, pH 7.5) at 16 °C. Currents from expressed channels 1-3 days post-injection were recorded in two-electrode voltage clamp (Oocyte Clamp OC-725C, Warner Instruments), digitized using a PowerLab 4/30 system (ADInstruments) and recorded using LabChart 8 software (ADInstruments). Data was obtained from at least two different batches of oocytes for each experimental group.

Oocytes were held at -70 mV and continuously perfused with perfusion buffer (2 mL/min). The perfusion buffer for oocytes expressing GABA_A_Rs was ND96 buffer (96 mM NaCl, 2 mM KCl, 1 mM CaCl_2_, 1 mM MgCl_2_, 5 mM HEPES, pH 7.5). The perfusion buffer for oocytes expressing Gly receptors was MBS buffer (88 mM NaCl, 1mM KCl, 2.4 mM NaHCO_3_, 19 mM HEPES, 0.82 mM MgSO_4_, 0.33 mM Ca(NO_3_)_2_, 0.91 mM CaCl_2_, pH 7.5). Drugs were diluted into the corresponding perfusion buffer on the day of the experiment using stock solutions of an appropriate concentration. Picrotoxin (PTX) was diluted from a 0.5 M stock solution in DMSO. GABA and Gly were diluted from 1 M stock solution in water. Propofol (PPF) was diluted from 30-100mM stock solution in DMSO. The recording protocol was as follows: a 10 s pulse of PTX was followed by a series of 20-40 s pulses of increasing concentrations of either GABA or Gly, followed by a co-application of GABA or Gly with propofol, and a final 10 s pulse of PTX. Pulses were sufficiently long to resolve the peak response, and inter-pulse intervals were 5-15 min to allow washout with buffer and currents to return to baseline. Current traces were individually detrended in MATLAB 2024b (Mathworks) by subtracting a spline fit to manually selected baseline regions in each trace. GABA or Gly concentration-response curves (CRCs) were fit with the Hill equation:

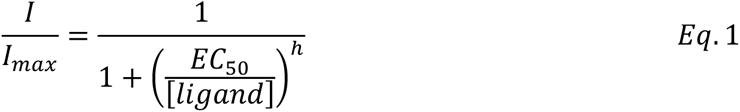

where 𝐼 is the magnitude of the ligand-elicited current, [𝑙𝑖𝑔𝑎𝑛𝑑] is GABA or Gly concentration, 𝐸𝐶_50_ is the concentration eliciting a half-maximal response, and ℎ is the Hill slope.

### Statistical analysis

Summary data was analyzed using Prism 10 (GraphPad). Symbols and error bars are mean ± SEM, and box plots show median and interquartile intervals. Where applicable, we applied Brown-Forsythe ANOVA, followed by Dunnett’s T3 multiple comparisons test. We focus on effect sizes that are at least multiple times larger than the background variation as opposed to relying solely on *P*-values.

### Molecular dynamics simulations

All-atom MD simulations starting from cryo-EM structures of closed (i.e., antagonist-bound, PDB 6X3S) or desensitized (i.e., GABA-bound, PDB 6X3Z) α1β2γ2 GABA_A_Rs in explicit solvent were conducted as described previously (Legesse et al., 2023). These structures, in the presence of relevant ligands, were placed in a simulation box with dimensions 127 × 127 × 163 Å^3^, embedded in a symmetric bilayer containing 40% cholesterol, 25% POPC (1-palmitoyl-2-oleoyl-sn-glycero-3-phosphocholine), 25% POPE (1-palmitoyl-2-oleoyl-sn-glycero-3-phosphoethanolamine), 9% POPS (1-palmitoyl-2-oleoyl-sn-glycero-3-phospho-l-serine), and 1% PtdIns(4,5)P2 (phosphatidylinositol 4,5-bisphosphate), previously shown to approximate the neuronal plasma membrane (Ingólfsson et al., 2017). Additionally, MD simulations of an advanced-sampled open state conformation for an α1β2γ2 GABA_A_R were conducted similarly to the closed and desensitized structures and as described previously (Haloi et al., 2025). For each structure, we simulated four 500 ns replicates using GROMACS-2024.2 or GROMACS-2025 (Páll et al., 2020).

We also carried out MD simulations of the ncAA amide-to-ester substitution β2(Leu140Lah) which we introduced into the desensitized structure (PDB 6X3Z). To achieve the β2(Leu140Lah) substitution, we replaced the pair of residues β2(Asp139-Leu140) with an Asp-like residue we termed Dap and the cognate α-hydroxy acid for leucine (Lah). Dap differs from Asp in that its carboxylate is covalently linked via an ester bond to the terminal hydroxyl oxygen of Lah. This swaps the amide peptide bond for an ester peptide bond without altering any sidechains. Noncanonical residues Dap and Lah were modeled within the CHARMM36m protein force field following the strategy used for backbone α-hydroxy and amide-to-ester substitutions in previous work (Infield et al., 2018; Leisle et al., 2022). In CHARMM topology space, the PRES EST patch converts a peptide bond to an ester by replacing the backbone amide N with an ester oxygen and using the standard CHARMM ester oxygen atom type OS together with the associated bond, angle, and dihedral parameters defined in the CHARMM36m parameter file. We adopted the same approach by adding Dap and Lah to the residue topology, hydrogen database, and residue type list using only existing CHARMM36 atom types, and the covalent ester linkage between the Dap sidechain carbonyl carbon and the Lah terminal oxygen was assigned bonded parameters similarly to the CHARMM36 ester terms used by PRES EST. Partial charges on Dap and Lah were chosen to preserve the intended net residue charge and to reproduce the CHARMM36 charge distribution of analogous peptide and ester groups as in previous work (Infield et al., 2018; Leisle et al., 2022). GROMACS reported no missing bonded or nonbonded parameters for Dap or Lah during preprocessing, and inspection of the full receptor simulations revealed no abnormal bond lengths or angles for the Dap–Lah linkage.

For simulations involving the ncAA Lah residue, we followed the same steps as for the closed and desensitized simulations. Briefly, the protein was embedded in a pre-equilibrated lipid bilayer generated with CHARMM-GUI using CHARMM36 lipid parameters. The final bilayer contained the same lipid composition as above arranged around the protein in a simulation box of the same dimensions. The system was solvated with TIP3P in a 150 mM NaCl solution to neutralize the system and approximate a physiological salt concentration. Ligand parameters were generated with the CHARMM general force field (CGenFF) to obtain CHARMM36-compatible GROMACS topologies.

The system was energy-minimized by undergoing sequential equilibration steps at 300°K with gradually released positional restraints to relax the system. The entire equilibration totaled 10 ns in duration. The first equilibration step involved a 2 ns NVT equilibration at 300°K with a V-rescale thermostat and three temperature coupling groups (Protein, Membrane, and Water + Ions) and included strong protein and ligand restraints (1000 kJ mol^-1^ nm^-2^ for all protein and ligand atoms). In the second 2 ns equilibration step, the backbone-only restraints were reduced to 100 kJ mol^-1^ nm^-2^ and the sidechain restraints were removed. In the final 6 ns equilibration step, all protein restraints were removed while retaining ligand restraints.

After energy minimization, the system was simulated for 500 ns in the NPT ensemble at 310°K to approximate physiological temperature. Temperature was controlled with a V-rescale thermostat, and pressure with a semi-isotropic Parrinello-Rahman barostat in both lateral and normal directions. A 2 fs time step was used in combination with LINCS constraints on all bonds involving hydrogens, which is standard for CHARMM36m and allows stable integration while maintaining accurate dynamics. Long range electrostatics were treated with a particle-mesh Ewald and van der Waals interactions were truncated with a force-switch between 1.0 and 1.2 nm in production. The Verlet cutoff scheme was used throughout.

### Phylogenetic analysis

Protein sequences of pLGICs from the organisms listed in **Supplementary Table 4** were retrieved from Uniprot proteomes based on the InterPro annotation as belonging to the “Neuronal acetylcholine receptor” (IPR038050) superfamily. Datasets per species were deduplicated with MMseq2 (Steinegger & Söding, 2018) and 95% sequence similarity. Deduplicated sequences were aligned using T-Coffee or MAFFT (SnapGene v8). Non-pLGIC sequences and fragments were confirmed using AlphaFold-generated structures and discarded. Sequences lacking the M2 α-helix and/or the M2-M3 linker were also discarded. The final alignment per organism was searched for anionic and cationic signature motifs at the beginning of the M2 α-helix: PXR, with X a non-charged residue, as the anionic motif, and E/D/Q followed by R/K/Q as the cationic motif. Sequences that failed to possess any of these motifs were discarded, except for proteins experimentally tested (mostly *C. elegans* subunits). The final 368 selected sequences were then aligned using MUSCLE (MEGA12) (Stecher et al., 2020, 2026) and the residue in the position of interest in M2-M3 (fourth residue following a conserved proline in the M2-M3 linker) was identified. The alignment was then used for tree construction and visualization using MEGA12. The phylogeny was inferred using the maximum likelihood method and Poisson model of amino acid substitutions (Zuckerkandl & Pauling, 1965). The tree with the highest loglikelihood is provided in Supplementary File 1 and at https://itol.embl.de/shared/1ieszf2NJt15U. The number of replicates (196) was determined adaptively. The percentage of replicate trees in which the associated taxa clustered together is shown next to the branches. The partial deletion option was applied to eliminate all residue positions with less than 90% coverage in the alignment, resulting in a final data set comprising 246 aligned positions.

## Acknowledgements

This research was supported by NIH grants R01GM148591 to M.P.G-O. and R35GM148239 to C.A.A., and by Swedish Research Council (VR) grants 2019-02433 and 2021-05806 to R.J.H. and E.L. Additional support was provided by the Knut & Alice Wallenberg Foundation and Swedish e-Science Research Center to R.J.H. and E.L.

## Contributions

M.P.G.-O. conceived and supervised this work. J.D.G. carried out, and C.A.A. supervised, the ncAA synthesis. C.M.B., M.P.G.-O., and C.A.A. established ncAA protocols for TEVC recordings. C.M.B., N.G.D., P.G. and S.K.S. carried out, and C.M.B. analyzed, the TEVC recordings. L.S. carried out and analyzed, and M.P.G.-O., N.H., R.J.H. and E.L. supervised the molecular dynamics simulations. C.M.B. carried out and H.H.Z. supervised the phylogenetic analysis. C.M.B., L.S. and M.P.G.-O. produced the visualizations. C.M.B., L.S. and M.P.G.-O. wrote the manuscript.

## Additional information

#Supplementary Information and a Supplementary File 1 are available for this paper. Correspondence and requests for materials should be addressed to Marcel P. Goldschen-Ohm.

### Supplementary File 1

Phylogenetic tree of pLGICs, identifiable as forming either anionic or cationic channels, from unicellular and multicellular eukaryotes. The tree was constructed using the maximum likelihood method, with the number of replicates (196) determined adaptively. The percentage of replicate trees in which the associated taxa clustered together is shown next to the branches. The bootstrap range values represent the last estimated bootstrap support value for that branch - the upper bound (or confidence limit) estimated during the adaptive bootstrap procedure. The branch labels indicate [UniProt ID] [organism code] [anion-/cation-selectivity] [M2-M3 residue at the homologous position with the conserved tyrosine]. The organism name codes can be found in Supplementary Table 4. The color of the lines indicates anion-/cation-selectivity and the M2-M3 residue (blue: Anionic/Tyrosine; violet: Anionic/Non-tyrosine; red: Cationic/Non-tyrosine; orange: Cationic/Tyrosine).

## Supplementary Information

**Supplementary Figure 1.**
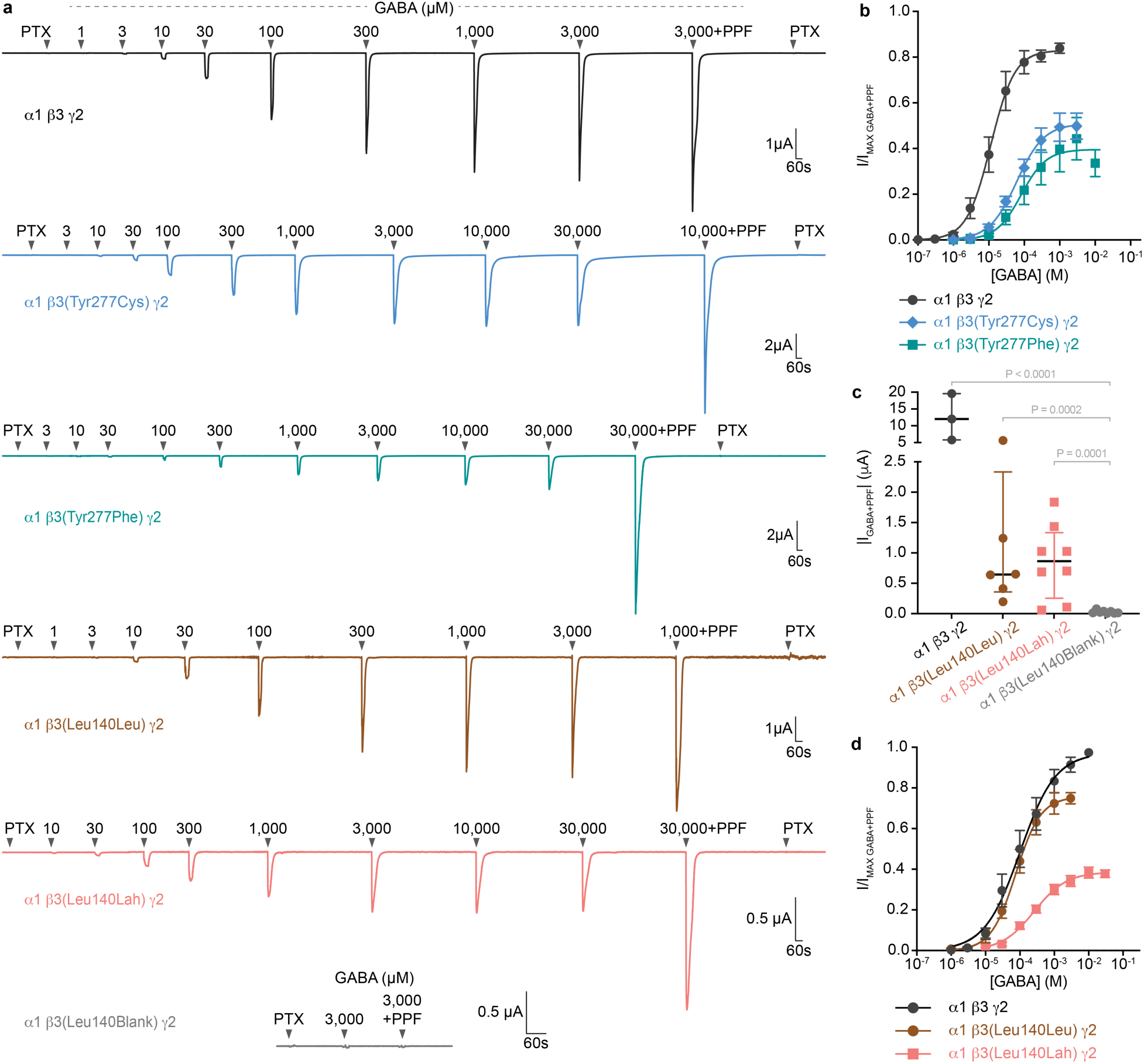
A H-bond linking gating loops is functionally important in GABA_A_ β subunits. **a** Representative current traces for α1β3γ2 GABA_A_Rs and either 1) substitutions of the conserved tyrosine β3(Tyr277) or 2) nonsense suppression incorporation of either wild-type leucine or its cognate α-hydroxy acid (Lah) at β3(Leu140). Arrows above traces indicate approximate onset of pulses of either 1 mM picrotoxin (PTX), increasing concentrations of GABA, or maximal GABA + 30 µM propofol (PPF). **b** Concentration-response relations for GABA-elicited currents, normalized to maximal GABA + PPF responses. Data are mean ± SEM across oocytes. Curves are fits of the Hill equation (Eq. 1) to the means. See Supplementary Table 3 for summary statistics and fit parameters. Number of oocytes (n) are: α1β3γ2; n = 6; α1β3(Tyr277Cys)γ2: n = 4; α1β3(Tyr277Phe)γ2: n = 6. **c** Total current per oocyte obtained in the presence of maximal GABA + 30 µM propofol suggests reliable nonsense suppression incorporation of Leu and Lah with little to no read-through (i.e., relative lack of current for Blank). Number of oocytes for each condition (left to right) are n = 3, 6, 8, 8. Plots show median and interquartile intervals. P-values are from Brown-Forsythe ANOVA with posthoc Dunnett’s T3 multiple comparisons test. **d** Concentration-response relations for GABA-elicited currents, normalized to maximal GABA + PPF responses. Data are mean ± SEM across oocytes. Curves are fits of the Hill equation (Eq. 1) to the means. See Supplementary Table 3 for summary statistics and fit parameters. Number of oocytes (n) are: α1β3γ2: n = 3; α1β3(Leu140Leu)γ2: n = 6; α1β3(Leu140Lah)γ2: n = 8.

**Supplementary Figure 2.**
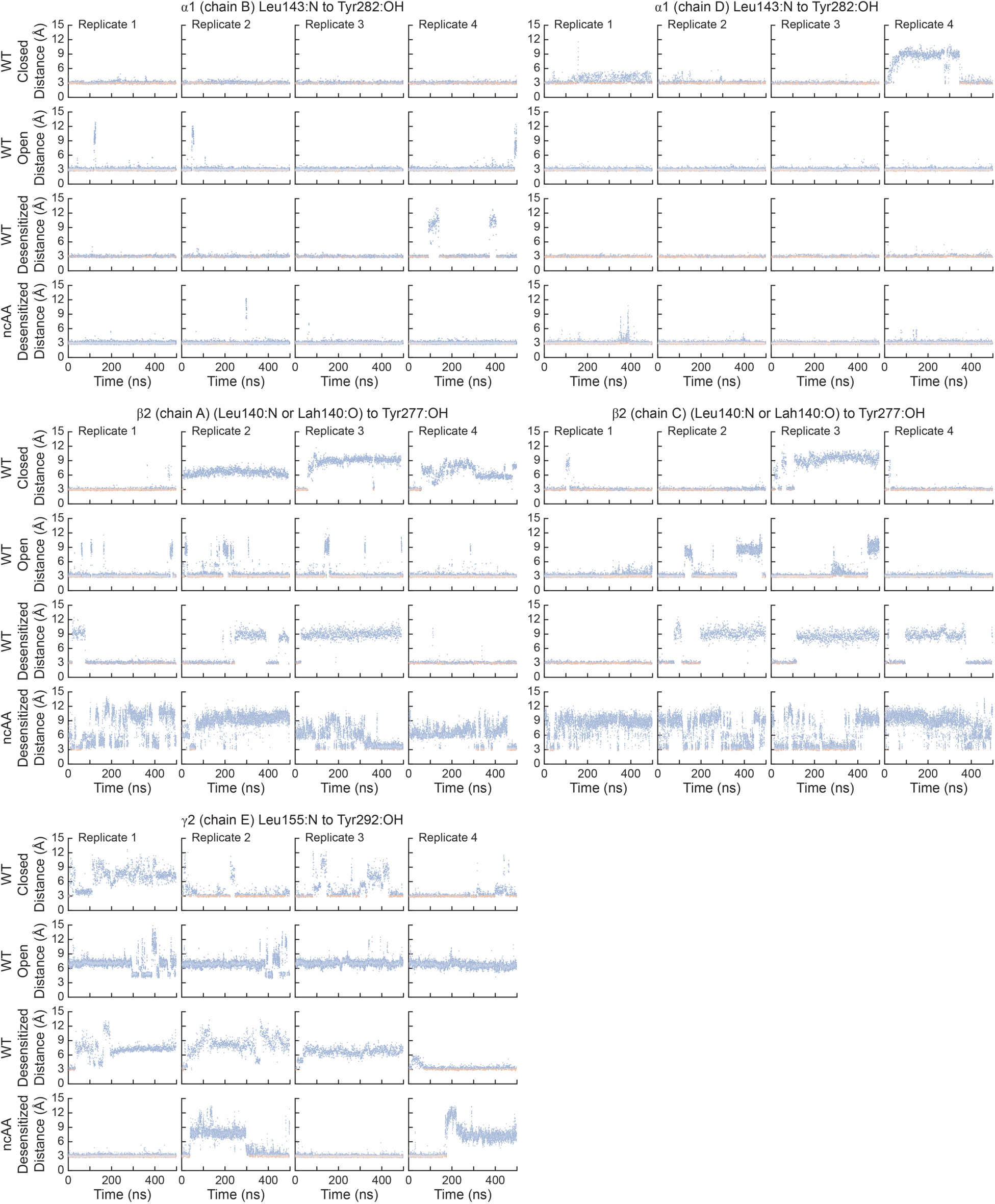
Time-resolved H-bond linkage between gating loops. Donor-acceptor distance trajectories from MD simulations of α1β2γ2 GABA_A_Rs in closed (antagonist-bound, PDB 6X3S), open (simulated structure (Haloi et al. 2025)), and desensitized (GABA-bound, PDB 6X3Z) states with and without the β2(Leu140Lah) amide-to-ester ncAA substitution. Distances are between a leucine backbone amide nitrogen in the Cys-loop (or the ester oxygen for the ncAA substitution) and a tyrosine OH group in the M2-M3 linker of the same subunit. Residue pairs are α1(Leu143:N-Tyr282:OH), β2(Leu140:N-Tyr277:OH), γ2(Leu155:N-Tyr292:OH), or β2(Lah140:O-Tyr277:OH). Data points are colored based on whether the atoms are within (red) or outside (blue) typical H-bond constraints (i.e., distance < 3.2 Å and bond angle in the range 120-150°).

**Supplementary Figure 3.**
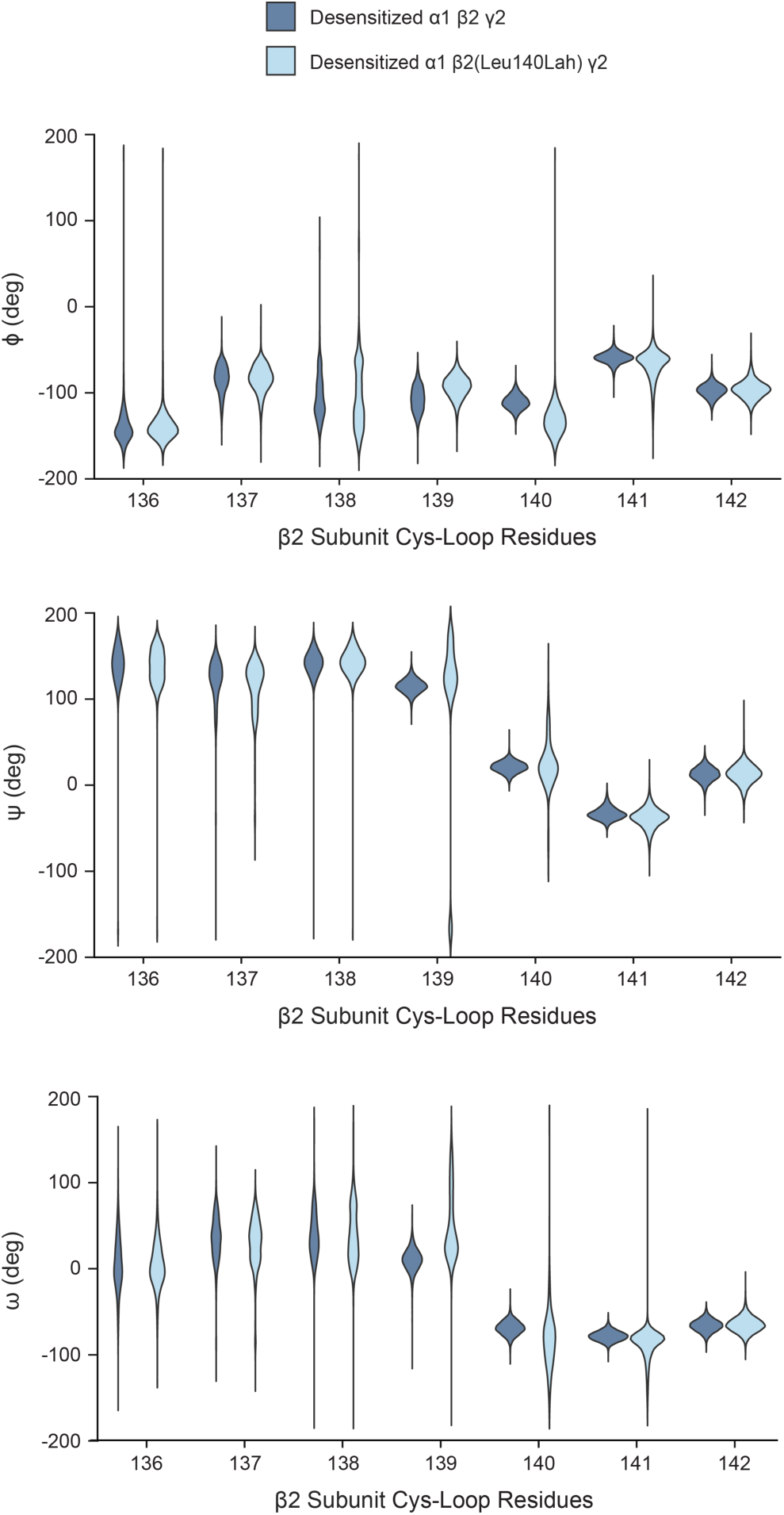
Increased backbone flexibility for an amide-to-ester swap that ablates a stabilizing H-bond. Backbone dihedral angle distributions ϕ, ψ, ω for Cys-loop residues 136-142 of β2 (chains A and C combined) from MD simulations of an α1β2γ2 GABA_A_R in a desensitized (i.e., GABA-bound, PDB 6X3Z) conformation with and without the β2(Leu140Lah) amide-to-ester ncAA substitution. ϕ is defined by the atoms *C_i-1_* – *N_i_* – *C*_α*i*_ – *C_i_.* ψ is defined by the atoms *N_i_* – *C*_α*i*_ – *C_i_ - N_i+1_.* ω is defined by the atoms *C*_α*i*_ – *C_i_* – *N_i+1_* – *C*_α*i+1*_. For β2(Leu140Lah), the backbone nitrogen is replaced by the corresponding ester oxygen in these definitions.

**Supplementary Figure 4.**
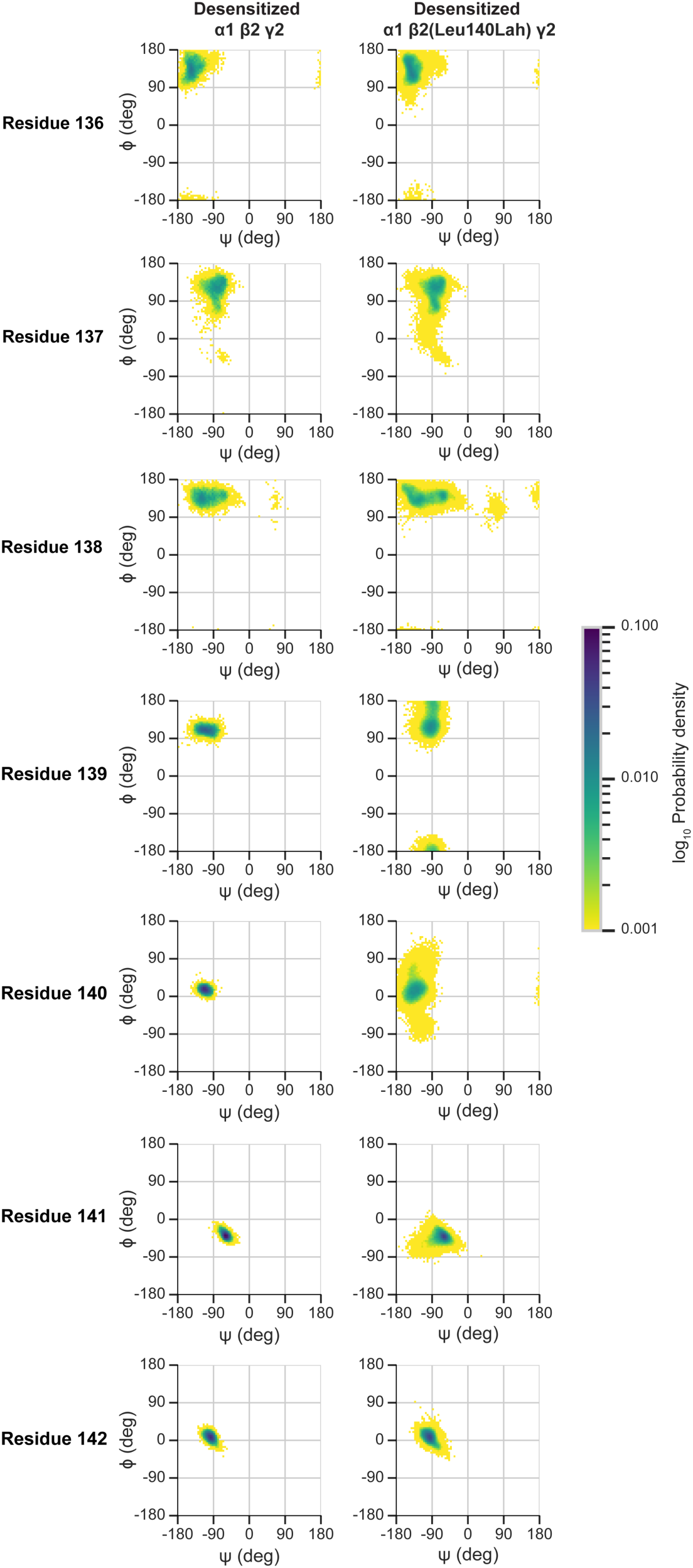
Increased backbone flexibility for an amide-to-ester swap that ablates a stabilizing H-bond. Ramachandran plots (Kleywegt and Jones 1996) of backbone dihedral angles ϕ and ψ for Cys-loop residues 136–142 of the β2 subunit (chains A and C combined) from MD simulations of an α1β2γ2 GABA_A_R in a desensitized (i.e., GABA-bound, PDB 6X3Z) conformation with and without the β2(Leu140Lah) amide-to-ester ncAA substitution. ϕ is defined by the atoms *C_i-1_* – *N_i_* – *C*_α*i*_ – *C_i_.* ψ is defined by the atoms *N_i_* – *C*_α*i*_ – *C_i_ - N_i+1_.* ω is defined by the atoms *C*_α*i*_ – *C_i_* – *N_i+1_* – *C*_α*i+1*_. For β2(Leu140Lah), the backbone nitrogen is replaced by the corresponding ester oxygen in these definitions.

**Supplementary Table 1.**
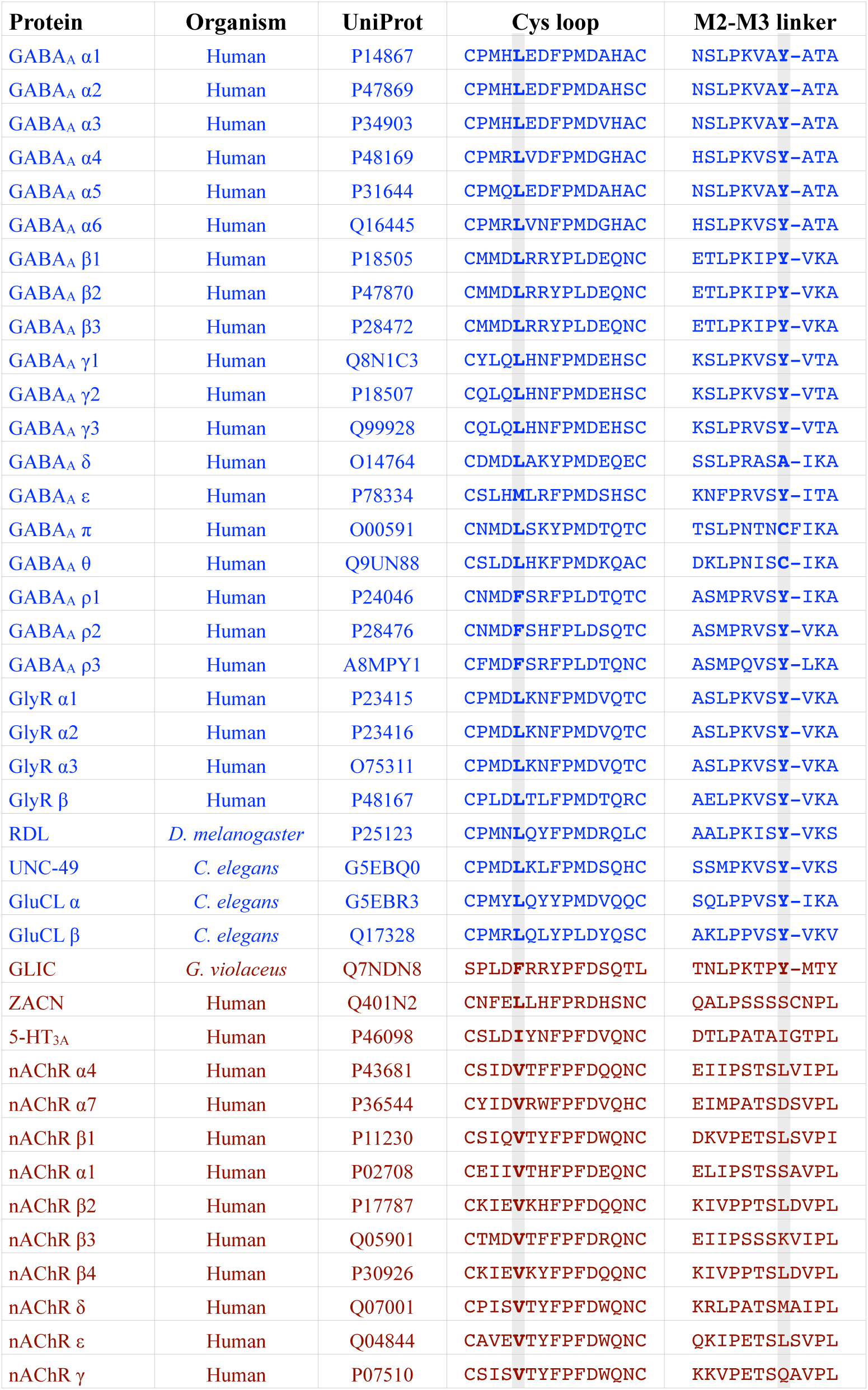
Sequence alignment of the Cys-loop and M2-M3 linker for inhibitory and excitatory pLGICs.

**Supplementary Table 2.** Variants in the M2-M3 position of inhibitory pLGICs subunits.

| Receptor | Subunit | Substitution | Mutation | ClinVar |  |
| --- | --- | --- | --- | --- | --- |
| GABA<sub>A</sub> | α1 | Y309>F | A>T | no report (COSV50102834) | Moderate Impact (predicted) |
|  | α2 | Y309>C | A>G | no report (NCI-TCGA: TCGA novel) | Probably damaging (predicted) |
|  | α2 | Y309>H | T>C | no report (COSV62915565) | no prediction |
|  | α3 | Y334>C | A>G | no report (NCI-TCGA: TCGA novel) | Moderate Impact (predicted) |
|  | α4 | Y315>D | T>G | no report (rs1184211379) | Probably damaging (predicted) |
|  | α5 | Y316>D | T>G | no report (COSV107422957) | no prediction |
|  | α5 | Y316>F | A>T | no report (COSV59476238) | no prediction |
|  | α5 | Y316>N | T>A | no report (rs1186567951) | Probably damaging (predicted) |
|  | α6 | Y299 none reported |  |  |  |
|  | β1 | Y302 none reported |  |  |  |
|  | β2 | Y301>C | A>G | RCV000479025, RCV000688521 | Pathogenic |
|  | β2 | Y301>F | A>T | RCV001390027, RCV004798911 | Pathogenic |
|  | β2 | Y301>H | T>C | RCV002035695 | Pathogenic |
|  | β3 | Y302>C | A>G | RCV001040961, RCV001311380, RCV003147577 | Pathogenic |
|  | γ1 | Y329 none reported |  |  |  |
|  | γ2 | Y331>C | A>G | RCV000998485, RCV001858879 | Pathogenic |
|  | γ2 | Y331>N | T>A | RCV002943951 | Uncertain significance |
|  | γ3 | Y312>C | A>G | no report (rs1891306645) | Probably damaging (predicted) |
|  | δ | A306 none reported |  |  |  |
|  | ε | Y336 none reported |  |  |  |
|  | π | C301>R | T>C | no report (rs1471896995) | Probably damaging (predicted) |
|  | π | C301>S | G>C | no report (rs756455222) | Probably damaging (predicted) |
|  | θ | C324>Y | G>A | no report (rs1556820226) | Benign (predicted) |
|  | ρ1 | Y340>C | A>G | no report (rs754011759) | Probably damaging (predicted) |
|  | ρ2 | Y320 none reported |  |  |  |
|  | ρ3 | Y326>H | T>C | no report (rs895150804) | Probably damaging (predicted) |
| Glycine | α1 | Y307>C | A>G | RCV000017441, RCV001376583, RCV005620337 | Pathogenic |
|  | α1 | Y307>S | A>C | RCV000031892 | Pathogenic |
|  | α2 | Y312 none reported |  |  |  |
|  | α3 | Y312>D | T>G | no report (rs560093755) | Probably damaging (predicted) |
|  | α3 | Y312>H | T>C | no report (rs560093755) | Probably damaging (predicted) |
|  | β | Y325>C | A>G | RCV003091751 | Uncertain significance |

**Supplementary Table 3.** Parameters for Hill equation fits to GABA concentration-response data.

| Receptor | EC<sub>50</sub> (μM) | n<sub>H</sub> | I<sub>max</sub> | n | Figure |
| --- | --- | --- | --- | --- | --- |
| GABA<sub>A</sub>R |  |  |  |  |  |
| α1β2γ2 | 26 (21 to 32) | 1.09 ± 0.09 | 0.828 ± 0.014 | 5 | Fig. 2b |
| α1β2(Tyr277Cys)γ2 | 159 (116 to 227) | 0.89 ± 0.09 | 0.147 ± 0.004 | 7 |  |
| α1β2γ2 | 95 (71 to 133) | 0.92 ± 0.08 | 0.910 ± 0.024 | 6 | Fig. 2c |
| α1(Tyr281Phe)β2γ2 | 42 (31 to 58) | 0.93 ± 0.09 | 0.935 ± 0.027 | 10 |  |
| α1β2(Tyr277Phe)γ2 | 816 (471 to 1,740) | 0.67 ± 0.08 | 0.334 ± 0.015 | 5 |  |
| α1β2γ2(Tyr292Phe) | 57 (46 to 73) | 1.01 ± 0.07 | 0.829 ± 0.019 | 5 |  |
| α1β2γ2 | 77 (63 to 95) | 1.02 ± 0.07 | 0.773 ± 0.015 | 6 | Fig. 3e |
| α1β2(Leu140Leu)γ2 | 62 (49 to 81) | 1.48 ± 0.14 | 0.704 ± 0.015 | 6 |  |
| α1β2(Leu140Lah)γ2 | 322 (208 to 517) | 0.84 ± 0.13 | 0.252 ± 0.008 | 7 |  |
| α1β3γ2 | 11 (10 to 12) | 1.28 ± 0.05 | 0.830 ± 0.008 | 5 | Suppl.<br>Fig. 1b |
| α1β3(Tyr277Cys)γ2 | 82 (45 to 154) | 1.23 ± 0.31 | 0.396 ± 0.023 | 6 |  |
| α1β3(Tyr277Phe)γ2 | 61 (53 to 70) | 1.12 ± 0.06 | 0.509 ± 0.007 | 4 |  |
| α1β3γ2 | 102 (72 to 153) | 0.84 ± 0.08 | 0.973 ± 0.030 | 3 | Suppl.<br>Fig. 1d |
| α1β3(Leu140Leu)γ2 | 76 (71 to 82) | 1.20 ± 0.03 | 0.757 ± 0.006 | 6 |  |
| α1β3(Leu140Lah)γ2 | 261 (203 to 344) | 0.93 ± 0.07 | 0.386 ± 0.008 | 8 |  |
| GlyR |  |  |  |  |  |
| α1 | 186 (155 to 224) | 1.70 ± 0.14 | 0.967 ± 0.020 | 6 | Fig. 5b |
| α1(Tyr279Phe) | 2,970 (1,790 to 6,980) | 1.18 ± 0.22 | 0.446 ± 0.023 | 6 |  |

Hill equation (Eq. 2) parameters are from fits to the normalized responses of the corresponding EC<sub>100</sub> agonist plus propofol. EC<sub>50</sub> is expressed as mean (95% coefficient interval), n<sub>H</sub> and I<sub>max</sub> are expressed as mean ± SEM, and n is the number of oocytes. There are multiple entries for wild type receptors, because they were run in parallel with the different mutants.

**Supplementary Table 4.** Organisms used for the phylogenetic analysis and number of corresponding protein sequences encoding pLGICs.

| Organism | Code | Total sequences | Total Anionic | Anionic with Tyr | Total Cationic | Cationic with Tyr |
| --- | --- | --- | --- | --- | --- | --- |
| *Amphimedon queenslandica* |  | **0** |  |  |  |  |
| *Mnemiopsis leidyi* |  | **0** |  |  |  |  |
| *Homo sapiens* | HUMAN | **43** | **23** | 20 | **20** | 0 |
| *Drosophila melanogaster* | DROME | **38** | **20** | 13 | **18** | 2 |
| *Caenorhabditis elegans* | CAEEL | **83** | **40** | 37 | **43** | 2 |
| *Nematostella vectensis* | NEMVE | **82** | **31** | 31 | **51** | 10 |
| *Hydra vulgaris* | HYDVU | **30** | **14** | 11 | **16** | 0 |
| *Trichoplax adhaerens* |  | **0** |  |  |  |  |
| *Salpingoeca rosetta* |  | **0** |  |  |  |  |
| *Monosiga brevicollis* | MONBE | **1** | **0** | 0 | **1** | 0 |
| *Micromonas pusilla* | MICPS | **2** | **1** | 0 | **1** | 0 |
| *Micromonas commoda* | MICCC | **1** | **1** | 0 | **0** | 0 |
| *Emiliania huxleyi* | EMIHU | **15** | **3** | 1 | **12** | 6 |
| *Prymnesium parvum* | PRYPA | **8** | **3** | 0 | **5** | 2 |
| *Fragilariopsis cylindrus* | FRCYL | **3** | **0** | 0 | **0** | 0 |
| *Pseudo-nitzschia multistriata* | PNMUL | **4** | **0** | 0 | **4** | 0 |
| *Pseudo-nitzschia australis* | PNAUS | **1** | **0** | 0 | **0** | 0 |
| *Cylindrotheca closterium* | CYCLO | **1** | **0** | 0 | **0** | 0 |
| *Seminavis robusta* | 9STRA | **9** | **1** | 0 | **8** | 0 |
| *Ditylum brightwellii* | DIBRG | **2** | **0** | 0 | **0** | 0 |
| *Aureococcus anophagefferens* | AURAN | **13** | **9** | 9 | **4** | 1 |
| *Pelagomonas calceolata* | PLCAL | **12** | **2** | 1 | **10** | 3 |
| *Euplotes crassus* | EUPCR | **2** | **2** | 0 | **0** | 0 |
| *Stylonychia lemnae* | STYLE | **2** | **1** | 0 | **1** | 0 |
| *Symbiodinium microadriaticum* | SYMMI | **10** | **3** | 2 | **7** | 5 |
| *Symbiodinium necroappetens* | 9DINO | **6** | **1** | 1 | **5** | 4 |

