## Supplementary figures and images for "An atomic interaction conserved for over 600 million years gates inhibitory neurotransmission"

### Supplementary File 1

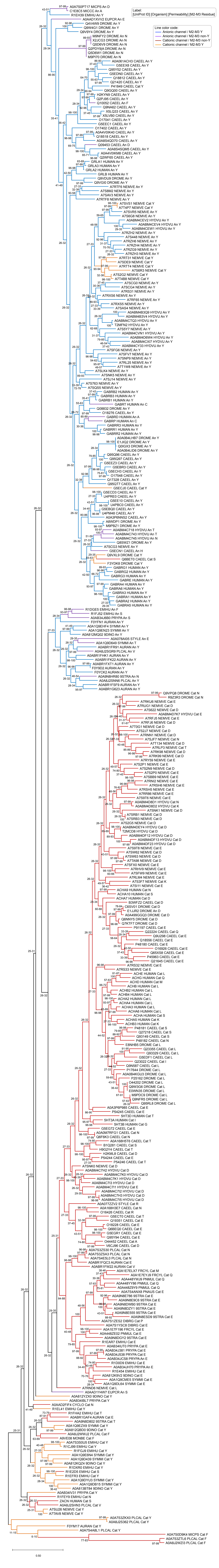
